# Physiological detection of flow states across varied gameplay workloads

**DOI:** 10.64898/2026.01.08.698463

**Authors:** James D’Amour, Reeshav Shrestha, Brent Miller, Brianna Croucher, Gaurav Sharma, Richard McKinley

**Affiliations:** Air Force Research Laboratory, Wright-Patterson AFB, USA; Henry M. Jackson Foundation for the Advancement of Military Medicine, Bethesda DC, USA; BAE Systems, London, United Kingdom

## Abstract

Flow is a cognitive state associated with heightened and seemingly effortless behavioral performance. Pioneering research suggests flow is an optimal cognitive mode for demanding tasks but the physiological underpinnings and identification on acute timescales, particularly across task contexts, remains an active area of research. Studies have demonstrated flow experiences are ubiquitous, in that they can arise during nearly any type of activity, suggesting flow represents a fundamental mechanism of skill mastery under which sensory inputs are seamlessly coupled to the necessary motor outputs. Cognitively, flow states are accompanied by a sense of complete task immersion, feelings of control, automatic behavior, decreased self-referential thinking and anxiety, changes in the perception of time, and improved affect to the point that arduous tasks can become autotelic, or self-rewarding. As wearable sensor technology continues to advance, the identification and manipulation of cognitive states in real-time is increasingly becoming a reality, with the potential to revolutionize learning, treatment of neurological disorders, and optimize task performance in domains ranging from professional sports to industry. This study aims to capture flow states from physiological measures so that they might be identified through wearable sensors coupled to machine-learning pipelines. In this work, models are made to predict high behavioral performance across disparately varied task conditions using multimodal physiological data, and these behavioral bouts are loosely correlated to participant responses to traditional flow state survey questions.

## Introduction

Flow is an optimal cognitive state for human performance associated with the subjective experience of total immersion in an activity (Csikszentmihalyi, 1975; Csikszentmihalyi, 1978, Csikszentmihalyi, 1990). Flow states are pervasive in culture, particularly in sports and other sensory-motor skills, where individuals may seem to effortlessly outperform competitors (Csikszentmihalyi, 1990; Jackson, 1996; Swann et al., 2016). However, these states are not limited to physical activities, as they have been documented in arithmetic, writing, art and other tasks (Katahira et al., 2018; Moneta and Csikszentmihalyi, 1990; Perry, 2009; Ulrich et al., 2014). Psychological research has associated flow states with increased drive, self-confidence, enjoyment, a greater sense of control, distortions in time perception, decreased self-awareness and abatement of anxiety, positioning them as a powerful mechanism to harness human drive, boost occupational fulfillment and the development of expertise (Csikszentmihalyi, 1975; Csikszentmihalyi and Csikszentmihalyi, 1990; Jackson and Edlund, 2004).

A sizeable body of work has documented the necessary, but not sufficient, conditions under which flow experiences may arise, mainly, clearly defined task goals, performance feedback and a match between participant skill and task difficulty (Csikszentmihalyi, 1975; Csikszentmihalyi and Csikszentmihalyi, 1990; Dietrich, 2004; Harris et al., 2017). However, a complete neurophysiological understanding of the flow state phenomenon remains an active area of research, with the potential to enable not only their detection with real-time wearables, but manipulation and induction (Ulirch et al., 2014; Gold and Ciociari, 2019).

Flow state research traditionally relies on Flow State Surveys (FSS) to document cognitive and affectual changes following stretches of task performance (Jackson and Eklund, 2004). Surveys broadly come with two inherent limitations. First, one must leave a potential flow state to reflect on if they were in a flow state. Second, as even abbreviated versions of these flow surveys contain five to twenty questions, they are difficult to incorporate into study designs aimed at capturing micro-flow states that might arise and dissipate on the scale of tens of seconds to a few minutes (or only a portion of the task block that the surveys are inquiring about). Wearable sensors coupled with algorithmic pipelines offer a greater level of temporal detail around the physiological signals associated with bouts of high-end human performance. However, a physiological approach is not without its own hurdles, mainly how generalizable are these markers between varying types of activities or are they capturing task-specific motor sequences? Additionally, within a given task how well can algorithms generalize between various difficulties or workload conditions, which are also known to alter many of the reported flow state biomarkers?

### Neurophysiological basis of flow experience

There is greater agreement around the peripheral physiological signatures of flow states. For more than a decade, groups have documented correlated changes in electrodermal measures, respiration, and heart rate variability during flow-like experiences and high-end task performance (Chanel et al., 2008, 2011; Chin and Kales, 2019; Harmat et al., 2015; Harris et al., 2017; Keller et al., 2011; Leger et al., 2014; Ulrich et al., 2016). Flow-related bouts seem to arise without the need for a linearly matched increase in effort on the part of the performant (Csikzentmihalyi, 1975; Khoshnoud et al., 2020). This has led some to suggest flow states represent a fundamental shift in brain network efficiency for externally oriented tasks, as individuals become acutely focused on their actions seemingly at the expense of self-awareness (Csikszentmihalyi, 1975; Dietrich, 2004; Simlesa et al., 2018; Tan et al., 2023; Weber et al., 2009). Two non-mutually exclusive hypotheses around cognitive states and the production of flow experiences are suggested by the field, first the

Hypofrontality Theory suggests that Prefrontal Cortex (PFC), which is associated with conscious effort, working memory and self-awareness, is suppressed during flow state experiences in favor of networks that are more implicit and automatic (Dietrich 2004; Harris et al., 2017; Tan et al., 2023). The Synchronization Theory of flow more directly implicates attention as the driving force in flow state establishment, as it enables coherent oscillations between reward and attentional networks (Posner et al., 1987; Weber et al., 2009). A reasonable interpretation of flow establishment might be that attentional mechanisms act to differentially gate and permit neuronal synchronization between the regional brain sub-networks suggested by the Hypofrontality model.

A common subjective observation of flow is an experience of decreased self-awareness (Sadlo 2016; Ulrich et al., 2016). Self-referential thinking and awareness are tightly linked to activity in a group of coactive structures termed Default Mode Network, which is mutually antagonistic with another group of structures, the Central Executive Network (Goldberg et al., 2006; Menon and Uddin, 2010; Ulrich et al., 2022). These two networks have anticorrelated activity such that elevated activity in the DMN occurs with decreased CEN activity. This feature is largely mediated by the interconnectivity provided by the Salience network (SN), crucial in the directing of attentional resources. This positions the structures of the SN as potential intervention targets for the control of flow states (Menon 2015; Menon and Uddin 2010; Sridharan et al., 2008). Interestingly, DMN network activity is chronically elevated among depressed individuals, and connectivity analysis has demonstrated strong interactions of this network with the subcortical structures of the amygdala and hippocampus that are involved in emotion valence and memory formation (Broyd et al.; 2009; Posner et al., 2016). It seems possible that depression effectively represents an anti-flow state, where-in the CEN that attends external tasks and stimuli, is dampened by continual loops of self-referential thinking, or rumination, that accompanies heightened DMN activity (Menon, 2019; Sadlo, 2016). Which may partially explain why exercise is effective in the treatment of depression, as it forces individuals to break DMN dominated brain activity and attend to external task challenges (Cline et al., 2024). As elements of these brain structures are plastic, continued retraining through habitual exercise can effectively normalize baseline network activity levels, alleviating depression and with the potential to be applied to several other disorders (Liang et al., 2015; Posner et al., 2016).

### Physiological markers of optimal experience

Experimentally flow literature has been mixed in support of the Hypofrontality Theory, as some groups have demonstrated results not necessarily in support of blanket frontal activity suppression during flow (De Sampaio Barros et al., 2018; Harmat et al., 2015; Yoshida et al., 2014). This suggests the need for greater resolution in dissection of neuronal structures that might better be mapped to networks like the CEN and DMN. As scalp EEG is temporally precise but spatially imprecise, this presents significant challenges to the identification of the relative activity levels of these subnetworks from wearables utilizing a limited number of dry-electrode recording locations. Harmat and colleagues employed functional near-infrared spectroscopy which can alleviate some of the challenges of EEG recordings, but did not detect region wide changes in frontal activity during flow (Harmat et al., 2015). While other groups have demonstrated differential activity within subregions of the PFC, specifically elevated activity within ventrolateral and dorsolateral prefrontal subregions, and suppression within the medial PFC (De Sampaio Barros et al., 2018; Yoshida et al., 2013). Functional magnetic resonance imaging studies have suggested increased activity in lateral PFC (inferior frontal gyrus) and posterior cortical regions with concurrent suppressed activity in medial PFC, the posterior cingulate cortex and amygdala, structures of the DMN (Ulrich et al., 2014). Although it remains possible that the structures involved in flow experience are somewhat task specific, a theme from much of this work is that flow is mediated by the more lateral CEN structures of the PFC (dorso and ventro-lateral PFC), while medial structures of the PFC tied to the DMN are suppressed (De Sampaio Barros et al., 2018; Ulrich et al., 2014; Ulrich et al., 2016; Yoshida et al., 2014). These regional differences in activity have proven a suitable target for flow state manipulation through transcranial direct-current stimulation (Gold and Ciorciari, 2019).

Most existing solutions to cognitive state decoding rely on features extracted from oscillatory power of typical neuronal frequency bands, spanning from 1-40 Hz. In previous work, Katahira and colleagues demonstrated increased theta activity in frontal areas, but as theta has been linked to working memory it is possible this is a result of increased workload and not flow experience per se (Katahira et al., 2018). Other work has suggested that alpha band power might be a better marker of flow-state onset particularly compared between hemispheres (Berta et al., 2013; Leger et al., 2014; Wolf et al., 2015). Promisingly, sensorimotor beta power has been shown to increase during fluid motor actions associated with flow performance and becomes synchronized across cortical regions as the subjective rating of flow experience increases, suggesting accurate oscillatory markers that differentiate between workload and flow might be derived from parietal beta to frontal alpha or theta ratios (De Kock, 2014; Koushnoud et al., 2020; Moreno et al., 2020). Decoupling these observed changes from various motor output requirements in differing tasks and varying task difficulties to extract common features that undergird the development of a true flow state remains a cutting-edge challenge in flow state detection research.

One way this can be addressed is using multimodal physiological sensing from disparate organs of the body by capturing peripheral measures of heart rate, respiration, ectodermal activity, and pupillary information along with extracted EEG features. Recent work modelling performance has shown promise with multimodal approaches, particularly with features derived from the eyes (Tashev et al., 2024). In this approach oscillatory band powers can be considered in the context of pupil size, or heart rate activity permitting higher resolution and separation of cognitive states.

## Results

### Behavioral analysis and flow surveys

To examine the physiological hallmarks of putative flow states during high behavioral performance, we collected scalp EEG, pupillometry, respiration, heart rate and survey data while participants played a custom video game task at varied difficulties, over three days. Study participants (n=19) were recruited according to the Air Force Research Lab (AFRL) IRB approved protocol, FWR20210142H (see methods for details). Within the experiment, participants play the Air Force Space Shooter Task (AFSS), in which a controller is used to navigate a ship in a two-dimensional space with asteroids continuously appearing from the sides of the screen (Fig 1A). Participants were instructed to destroy as many asteroids as possible without dying, which occurs when players collided with an asteroid or were killed by enemy (computer-controlled) ships. Ship deaths are not directly penalized in the displayed score but take time to respawn the player’s ship in the middle of the screen representing a task reset and direct feedback event of performance. This sensory motor task design is suitable for measuring continuous performance and attention, has clear goals, immediate sensory feedback through visual and auditory modalities, and provides a straightforward way of scaling difficulty for the experimenter. In the AFSS, the participant’s navigable space on screen is inversely related to the number of asteroids presented and destroyed asteroids are immediately replaced. Task difficulty is manipulated by changing the number of asteroids present. More asteroids make participants increasingly rely on destroying or navigating around obstacles to keep the ship alive.

**Fig. 1.**
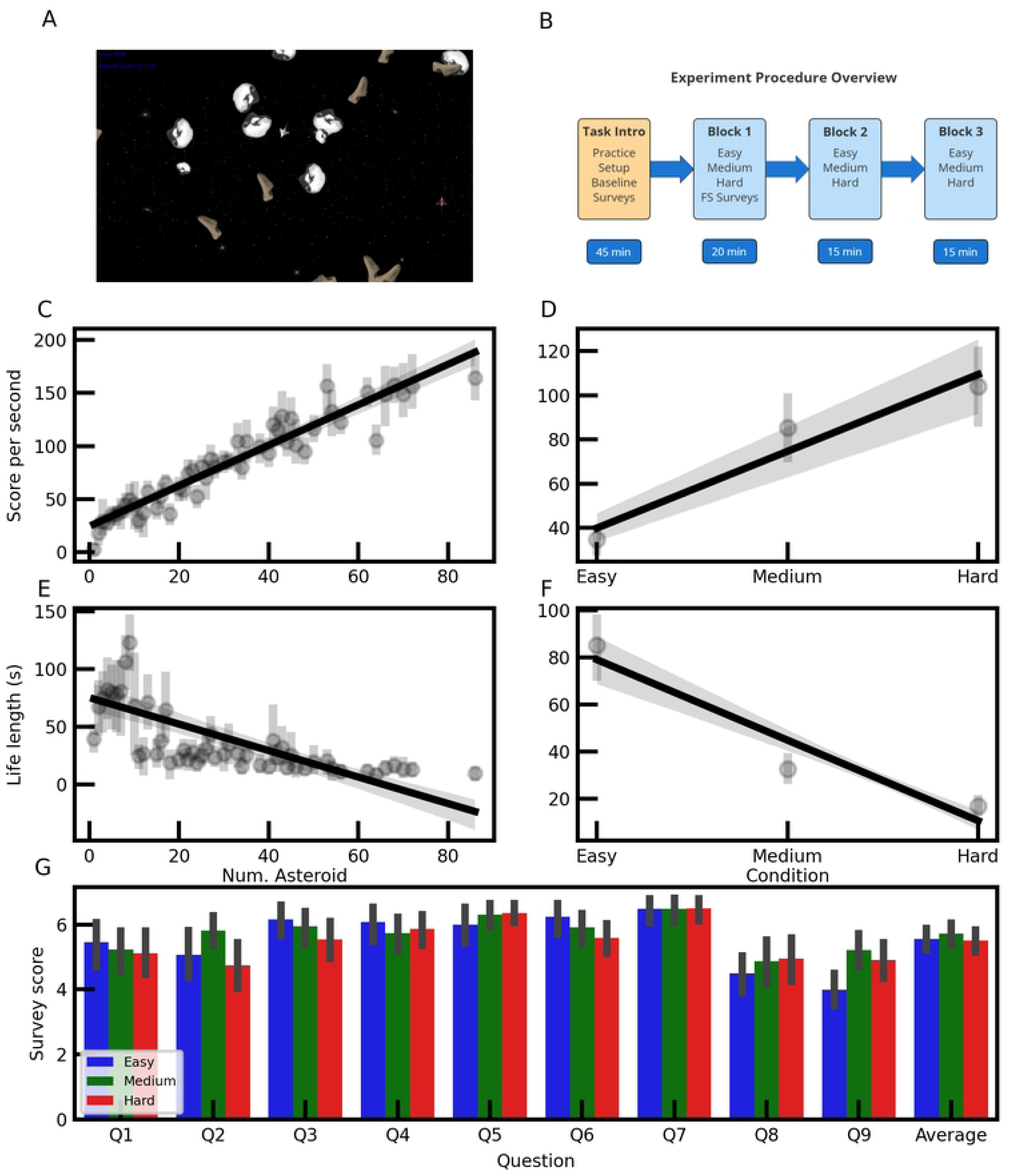
Behavioral quantification across task difficulty. (A) Gameplay image of the AFSS task, (B) Experimental task design, (C) Participant score per second against asteroid number, r = 0.95, slope = 1.97 ± 0.09, p <0.001, (D) Same as in C, but with normalized workload conditions, r = 0.69, slope = 34.55 ± 4.91, p < 0.001. (E) Participant life length against asteroid number, r = −0.71, slope = −1.20 ± 0.16, p < 0.001, (F) Same as in E, but with normalized workload conditions, r = - 0.83, slope = −34.14 ± 3.06, p < 0.001. (G) Difficulty block averaged flow state survey responses, F = 2.20, p=0.13, Fig 1G, right most bars; E 5.56 ± 0.12, M 5.72 ± 0.12, H 5.51 ± 0.12, avg and sem reported.

Importantly the design permits participants to employ a broad range of behavioral strategies to achieve success, as long as they maximize score and minimize deaths. Similar video-game based experimental designs have been used previously in physiological flow state research, with groups typically inferring that high performance is associated with heightened flow experiences, though it is unclear if flow induces high performance, or high performance induces flow experiences (De Kock, 2014; Khoshnoud et al., 2020; Nunez-Castellar et al., 2016). In this work we take a behavior driven approach to detect potential flow experiences, and did not detect differences in flow state survey measures when collapsed across task difficulty alone (Fig 1G).

For each of the three days in lab, baseline participant skill was assessed, and normalized task difficulty levels of easy, medium and hard (E, M and H, respectively) were established by increasing or decreasing asteroid numbers relative to their baseline skill level (Fig 1B; see methods for full description). Participants face five-minute blocks of each custom-scaled difficulty level in a pseudo randomized order, that is repeated three times per day in the laboratory. The in-game score (displayed on screen to participants, top left corner Fig 1A) is derived from the number of asteroids destroyed during that difficulty block. Scores from destroyed asteroids (game-score) were highly correlated to objective task difficulty (actual asteroid number) and subjective difficulty (normalized to skill levels), which was unsurprising as more asteroids meant more potential targets (Fig 1C and D). These linear regressions are performed on participant average asteroid level and participant condition collapsed averages, respectively, and may differ slightly from the regression shown for display purposes in the figure. In comparing Figs. 1C and 1D the effect of normalizing task difficulty compared to raw asteroid number appears to be a decrease in the quality of the linear fit, suggesting a plateauing effect of task behavior is more readily observed under normalized task difficulties, particularly when aggregating across individuals.

Examining time survived revealed a negative correlation with difficulty; fewer asteroids were associated with longer life-lengths (Fig 1E and F). We noted that asteroid number (objective difficulty) was less strongly correlated with life-length than with game-score (absolute value of correlation, Fig 1C vs E). However, the inverse was true when comparing workload-condition (subjective difficulty) to life-length and game-score (Fig 1D vs F). Importantly, neither behavioral metric from gameplay displayed a peak within the skill-matched medium difficulty level, making them ill-suited for generalized flow state detection across workload conditions.

An abbreviated version of the Flow State Survey was administered after the first block of each difficulty level, on each day (Fig 1B and G; see supplemental S1). An ANOVA on the average survey responses across conditions failed to detect any significant differences. But some individual questions did trend toward significance between at least one workload level: Question 2 asked participants to rate their agreement with the statement, “Task demands were well matched to my ability”. Differences were detected between the M and H workload conditions (p-adj = 0.003), question 6, “I felt in total control of my actions”, showed differences between the E and H conditions (p-adj = 0.019), and question 9, “I was thrilled”, showed differences between both E - H and E - M conditions (p-adj = 0.004, <0.001, respectively). Note that these surveys are given at the end of five minute blocks, whereas plotted data are from individual trials within blocks (15 trials per block) meaning there is an inherent difference in the temporal resolution of the survey and plotted epoch level data, where participants might be averaging their recollection of their gameplay for the past few minutes, or more likely biased by the recency effect (Ebbinghaus, H. 1913).

### Workload-specific models of task performance

Across participant machine learning algorithms were deployed against both behavioral metrics separately (game-score and life-length) within a single normalized workload condition (Easy, Medium and Hard; Fig 2A). These permitted the examination of separability for each behavioral variable and the associated physiological features selected by the models, without the confounding factor of task difficulty variation. Between both task metrics we found that life-length consistently produced higher Roc-auc binary classification scores when dividing each metric into high and low classes based on the group mean. Roc-auc curves for models trained and tested on unseen data from a single normalized workload condition showed the best performance in the H condition (Fig 2A). To address the question of how task workload affects model performance, we tested the best performing H model (logistic regression, without retraining) on test data from the E and M workload conditions (Fig 2B). In averaging life-lengths from participant epochs, based on the grouping assignment from the H model, results showed that the hard-trained model was able to successfully select longer life length epochs in the medium task-workload test data, comparable to the hard test data under which it was trained. In contrast, the hard-trained model failed to differentiate epoch life-lengths in the easy test data, suggesting that creating a model capable of generalizing across task-workload will face significant challenges when task-workload levels are disparately varied. This inability of the H-model to differentiate epoch life-lengths in the E-workload condition is a common challenge faced in behavioral quantification, particularly as it relates to flow, namely, how to scale and quantify task performance across varied task difficulties. While individuals lived longer in the easy condition and scored higher in the hard condition, these metrics fall short of capturing actual participant performance and are largely the result of the task features (more or fewer asteroids) as we know empirically that real participant skill levels are more closely matched to the medium difficulty level that was calibrated each day.

**Fig. 2.**
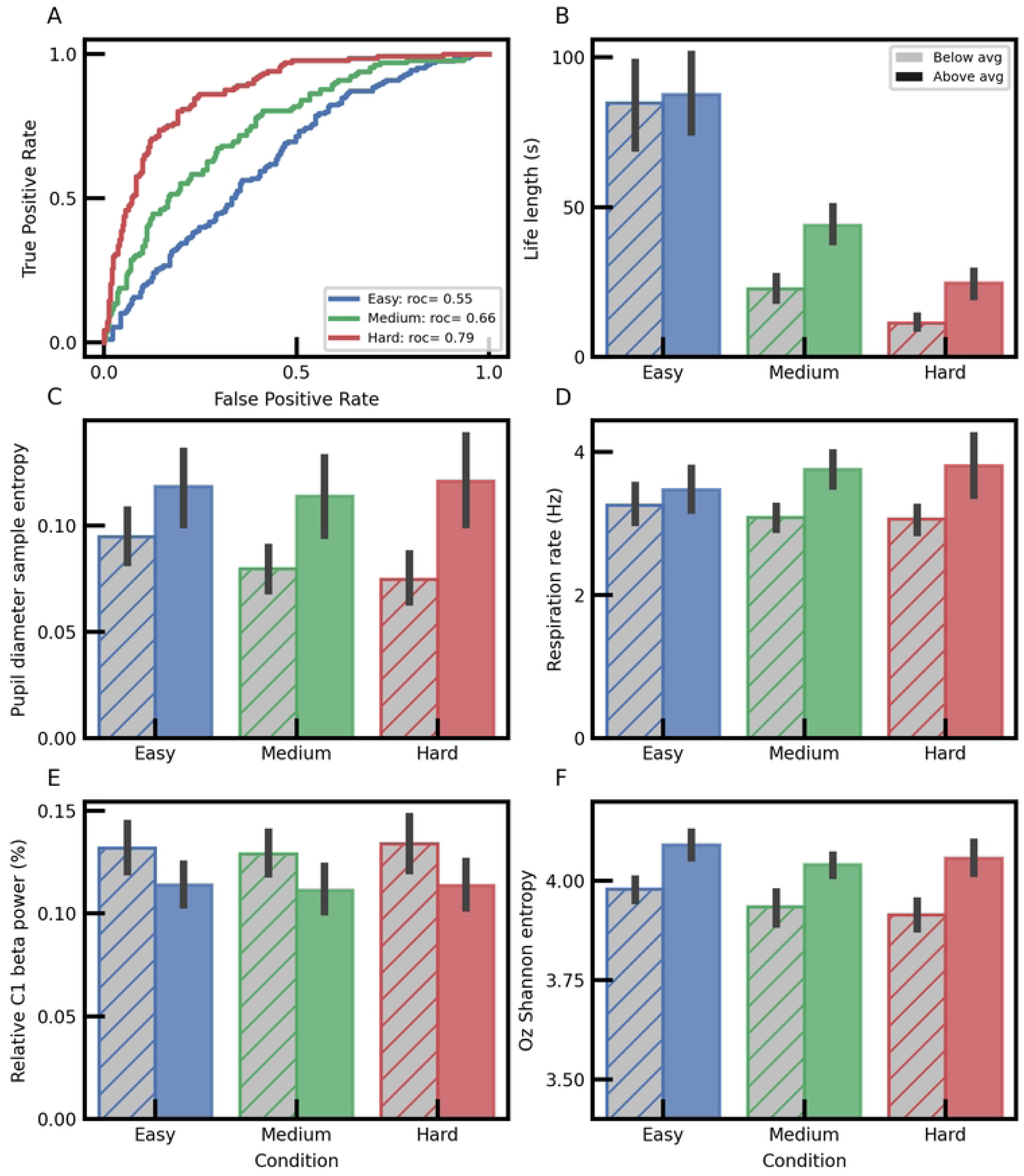
Physiological differences in binary classification models. (A) Roc-auc curves for binary classification using logistic regression models, trained and tested on unseen data from a single normalized task difficulty, 55%, 66%, 79%, for E, M and H, respectively. (B) Participant averaged life lengths from epochs assigned to low and high class labels using the best performing Hard model in A, ANOVA: F = 52.38, p < 0.001; post-hoc tests: life length values for predicted low and high class labels in H: 11.27 ± 1.01 vs 24.60 ± 2.18 seconds, p < 0.001; M: 22.75 ± 2.06 vs 43.95 ± 3.12 seconds, p < 0.001; E: 84.71 ± 7.70 vs 87.48 ± 6.64 seconds, p = 0.79. (C) As in B, but with pupil diameter sample entropy, ANOVA: F = 6.13, p-adj < 0.001; pupil diameter entropy values for predicted low and high class labels in H, respectively: 0.0747 ± 0.006 vs 0.12 ± 0.11 a.u., p-adj = 0.002; M: 0.08 ± 0.005 vs 0.11 ± 0.0097 a.u., p = 0.042; E: 0.95 ± 0.007 vs 0.118 ± 0.009 a.u., p-adj = 0.314. (D) As in B, but with respiration rates, ANOVA: F = 5.26, p-adj < 0.001; respiration rates for predicted low and high class labels in H, respectively: 3.06 ± 0.089 vs 3.80 ± 0.230 breathes per ten sec, p-adj = 0.005; M: 3.09 ± 0.087 vs 3.75 ± 0.126 breathes per ten sec, p-adj = 0.014; E: 3.26 ± 0.0128 vs 3.471 ± 0.150 breathes per ten sec, p-adj =0.870. (E) As in B, but with relative beta power from electrode C1 (%), ANOVA: F = 3.11, p-adj = 0.012; relative C1 beta band power values for predicted low and high class labels in H, respectively: 0.13 ± 0.006 vs 0.11 ± 0.006 a.u., p-adj =0.164; M: 0.13 ± 0.005 vs 0.11 ± 0.006 a.u., p-adj =0.278; E: 0.13 ± 0.006 vs 0.11 ± 0.005 a.u., p =0.256. (F) As in B, but with electrode Oz Shannon entropy, ANOVA: F = 16.013, p-adj < 0.001; Oz Shannon entropy for predicted low and high class labels in H: 3.915 ± 0.0189 vs 4.057 ± 0.021 a.u., p-adj < 0.001; M: 3.934 ± 0.021 vs 4.040 ± 0.0135 a.u., p-adj < 0.001; E: 3.979 ± 0.0137 vs 4.091 ± 0.0167 a.u., p-adj < 0.001.

Several of the most salient physiological features used in the model are shown at the participant averaged level (Fig 2). The participant-averaged epochs, selected by the hard model in both the medium and easy conditions are potentially informative as to feature utility. These all vary in magnitude in their flow-like and non-flow-like assigned epochs, but the nature of their difference over work-load condition reveals if that feature also displays sensitivity to task difficulty. For example, pupil diameter sample entropy diverges between the two performance groups as difficulty increases, whereas other features appear relatively insensitive to task difficulty level (Fig 2C-F).

### Participant and workload generalized models of task performance

A challenge faced by the field of flow-state modelling is the ability to function stably across varying task workloads, as changing task difficulty fundamentally shifts the distribution of task related performance metrics and some physiological features more tightly correlated with workload than performance (see Fig 1 C-F). In the AFSS task, harder difficulties mean higher game-scores (asteroids destroyed) and lower life lengths. These game play metrics are predictable within a task difficulty context (Fig 2), but models for generalized predictions across task difficulty are severely hindered by the overall shift in workload conditions (Fig 2B). For example, training models to predict life-length across all conditions essentially teaches the model to detect the easy trials, as participants nearly always lived longer in the easy condition. The problem is further exacerbated by the fact that the life-length variable would seem to encode different meaning at different workload levels-as suggested by the generalizability of the hard life-length model when tested in the medium and easy conditions in Figure 2. This makes some sense intuitively, as in sufficiently difficult task conditions life-length is a meaningful measure of task skill as one must be actively engaged in successful gameplay to keep their ship alive. However, in the easy condition life length becomes ambiguous – a long life here could signify high skill and engagement, or simply long periods of time without encountering many, if any, asteroid objects. Likewise, the inverse is true of the score metric.

We wondered if we could combine these two-task metrics to produce a generalized metric of performance that could function across workload context and be consistent with psychological flow literature expecting peak performance under participant-matched task difficulties. To do so, we placed a hard cap on life-length values (at 35 seconds) near the average value for life-lengths in the medium condition, but well below the average under the easy condition (Fig 1C and Fig 3A). As expected, this manipulation mainly affected easy condition epochs, but did place caps on some epochs in both other workload-conditions. Each epoch’s capped life-length value was then multiplied by each epoch’s game-score value resulting in a new metric, which we term performance. While the choice of capping life-length is somewhat arbitrary, this is an appealing approach as it allows our performance metric to recapitulate the expected inverted parabola shape of the Yerkes-Dodson law relating task-difficulty to optimal human behavior and the ‘flow-channel’ described in psychological literature as arising when challenge is closely calibrated to skill level (Fig 3B, Csikszentmihalyi and Csikszentmihalyi, 1990; Yerkes and Dodson, 1908). By using a binary cutoff based on the cross task-workload performance average, fewer putative flow-like epochs would be predicted from the easy and hard conditions, but due to the spread of the data some will still be included. This matches our understanding of cognitive states under workload conditions: participant matched workload (medium condition) is expected to be the most predisposed to the induction of flow states, but flow can still arise under easy and hard task conditions, though less often. Workload normalization was also crucial to constructing this metric, as the performance plotted against raw difficulty (asteroid number) was substantially less clear in distribution (Fig 3C). Though surveys were unsuccessful in showing condition dependent changes in Likert ratings between conditions (Fig 1G), we found that several questions were weakly positively correlated with this new performance metric (Fig 3D).

**Fig. 3.**
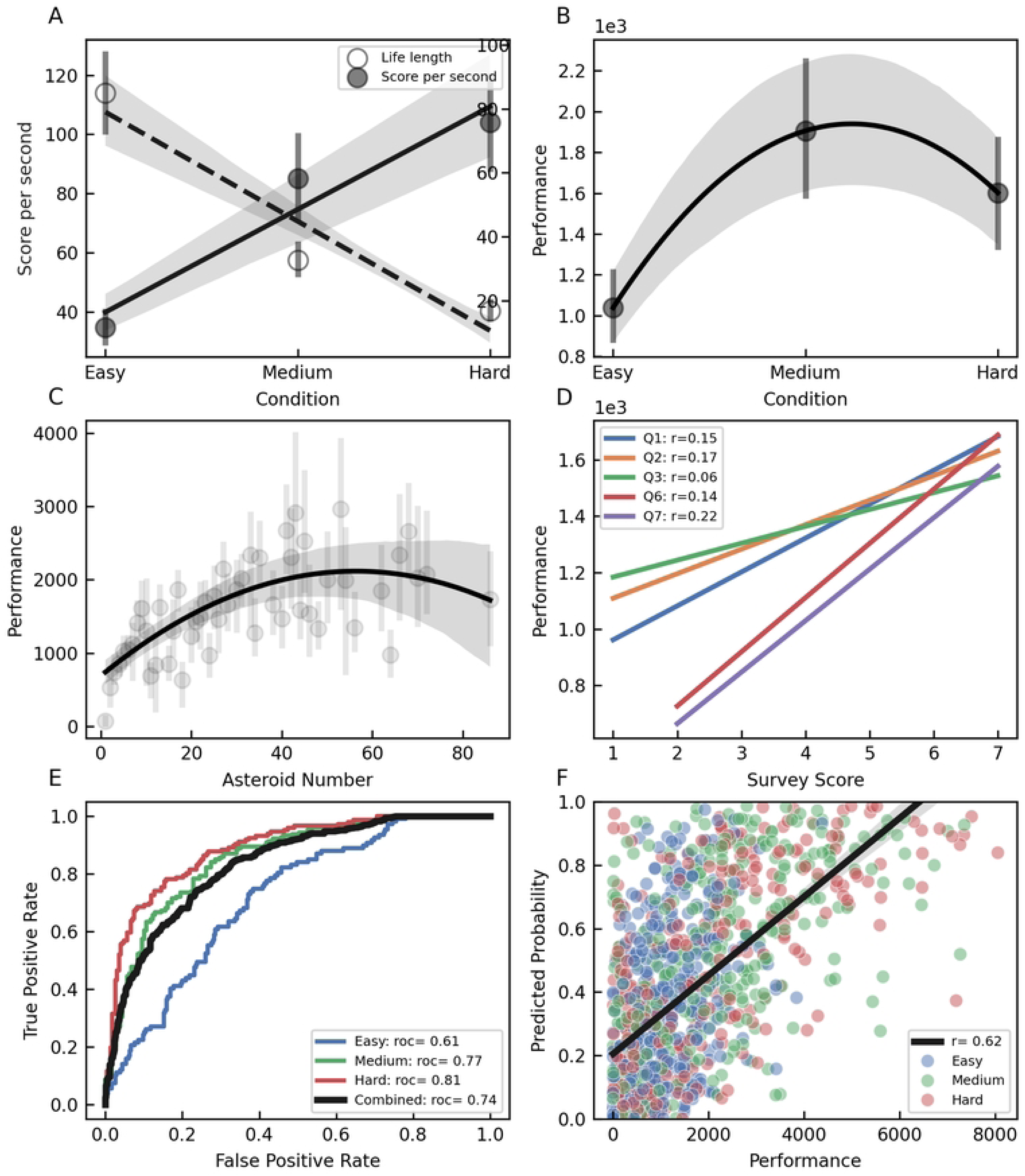
Performance classification across task difficulties. (A) Overlaid plots for score per second and life-length versus normalized task condition, same as in Fig 1 D and F. (B) Performance metric created by capping life lengths and multiplying by score per second, 1355 epochs were capped with a raw mean of 85.05 secs and a capped mean of 29.13 secs, 615 epochs capped with a raw mean of 32.58 secs and a capped mean of 21.17 secs, and 206 epochs capped with a raw mean of 16.83 and capped mean of 14.55 for E, M, and H, respectively. (C) Performance metric plotted against asteroid number. (D) Individual survey question responses plotted against averaged performance scores, with correlation values shown. (E) Roc-auc plots for logistic regression models tested on unseen data from all conditions, 74% overall. (F) Correlation of the predicted probability value from the combined model in E plotted against actual performance, r = 0.62, slope = 0.000124, p < 0.001.

Supervised machine-learning algorithms were trained and then tested against unseen data in a binary classification of high versus low performance (Fig 3E). These models were trained across participants and workload level. A logistic regression model showed a 74% Roc-auc score over all conditions for all participants, while deploying this model against only test data from each individual condition showed 61%, 77% and 81% Roc-auc scores for E, M and H conditions, respectively. Test data are stratified when being split out before training, so targets remain balanced across conditions. This model was used to generate probability distributions for each epoch, and these are shown plotted against actual performance (Fig 3F).

As the generalized model shown in Fig 3 contained greater than 900 features, we next examined feature importance with the aim of creating a reduced feature set model that retained classification accuracy (Fig 4). Recursive feature elimination showed that model performance plateaued at around 75 features (Fig 4A). Retraining the generalized model using these 75 top features and testing on unseen data resulted in a combined performance of 73% (Fig 4B). The top 25 features used in this model are shown in Fig 4C with their relative importance values. This model relied on some overlapping, but many differing features from those used by a single normalized workload condition model trained on hard data in Fig 2 (Fig 4 C versus D). Most notably, the generalized model utilized gamma from frontal, central and parietal electrode locations, whereas eye features seemed to play a more significant role in the hard-workload model. This suggests the possibility of multi-step classification pipelines as an effective approach to improved or tailored model predictions, as pupil related signals could be used to gate predictions generated from the model. We also noted that midline electrodes were largely absent from the top ranked feature sets. Instead most electrodes were laterally biased particularly in the frontal and central regions, suggesting that further feature engineering might be applied to these channels to create stronger flow-like physiological signals and that many of the other 64 recording sites might be largely redundant. It also agrees with the notion of laterality in flow states suggested by a growing body of literature (Gold and Ciorciari, 2016; Sridharan et al., 2008; Ulrich et al., 2022). In the workload specific hard model, pupil diameter sample entropy as well as the evoked pupil peak in response to the oddball tone were key features, where-as in the generalized model resting pupil diameter appeared more informative with the other ocular features being relatively less important. Heart rate was a prominent feature in both models. Respiration was less crucial in the generalized model than the hard model which could suggest it has greater sensitivity to workload condition, however we did not extensively derive features from the respiration, as was done with some other signal sources.

**Fig. 4.**
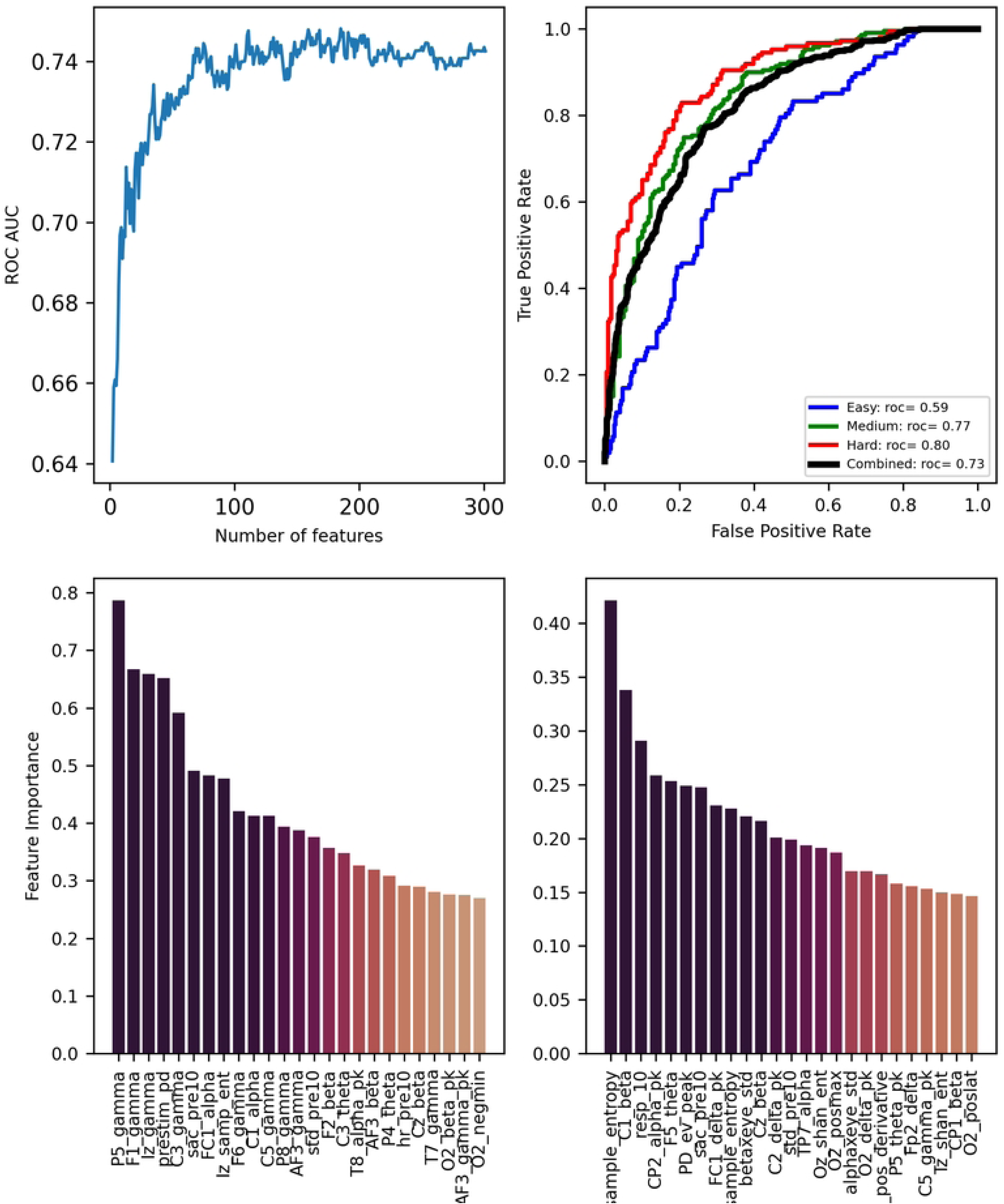
Reduced feature set model for generalized flow state prediction. (A) Recursive feature elimination displaying feature number and Roc-auc scores. Model performance plateaus near 75 features. (B) Roc-auc plot for the generalized reduced feature model tested on unseen data. (C) Top 25 features plotted against relative importance for the reduced generalized model. (D) As in C, for the hard workload-specific model created in Fig 2. For features in C and D, electrode location and a derived feature are separated by an underscore, for instance P5_gamma is the relative gamma power from parietal electrode 5 in the 10-20 layout. Any electrode bandpower followed by ‘pk’ refers to peak spectral power over the prestimulation interval, ‘negmin’ refers to the negative peak of the EEG event following the event, ‘posmax’ refers to the positive EEG deflection and ‘poslat’ is the associated latency of this peak. In all features ‘samp_ent’ and ‘shan_ent’ refer to the calculated sample entropy and Shannon entropy of the respective signal. ‘Prestim_pd’ is average pupil diameter in the ten seconds prior to event trigger, while ‘pd_sample_entropy’ refers to the sample entropy calculated from that same data segment. ‘PD_ev_peak’ is the positive peak of the pupil diameter following the event. ‘Sac_pre10’ is the saccades detected in the ten seconds prior to the event. ‘Samp_ent’ refers to the calculated sample entropy prior to event trigger. ‘Std_pre10’ is the standard deviation of the pupil in the ten seconds prior to the event trigger. ‘Resp_10’ is the breathing rate prior to the event. ‘Hr_pre10’ is the heart rate. ‘Betaxeye_std’ is the beta power from electrode CPz multiplied by the standard deviation of the pupil. ‘Alphaxeye_std’ is the same, except with alpha bandpower from electrode CP3.

It is likely that actual flow-states represent only a portion of the collected dataset and assuming half of them are ‘flow’ is likely an oversimplification. However, this approach allows us to screen and select for physiological signals tied to specific sensors, suggesting which sensors might be necessary and crucial in flow state detection and how to make them robust to changes in workload. Future work might benefit from unsupervised clustering approaches to identify putative flow epochs and creating models with unbalanced target classes, but even here assumptions might be made about the potential of every individual to produce a flow state.

## Discussion

The present work furthers the development of wearable sensor solutions for the detection of optimal cognitive states, specifically flow. This study took a multi-modal approach based on brain, eye, and other peripheral signals to create flow-predictive machine learning models that are robust to changes in task workload. This is a crucial step in the development of software for the emerging market of commercially available wearable sensor devices, as the ability to generalize across not only participants, but task contexts is likely to be a limiting factor to mass adoption and real-world utility. As such, the features and models discussed above provide a framework for the creation of cognitive state detection software that is task context agnostic. These models are not meant to be fully optimized and we suspect that individualized models and further feature engineering could produce still better model performance. Instead, this work explores the signal space and relative importance to decoding of high-performance cognitive states. Validation of these models in different visuo-motor video-game tasks would further advance and solidify this work, while contrasting it with auditory or other sensory dependent tasks might reveal interesting findings such as less reliance on ocular features or a differing set of EEG electrodes for model predictions.

There are notable differences between the feature set for models created within a workload-context (Fig 2), and models optimized for prediction across workloads (Fig 3 and 4), that likely carry important scientific implications and neural insights, such as the relative importance of EEG derived gamma powers in the latter. At the very least, it highlights the sensitivity of some of these features to both workload and performance, shedding some light on the parameter space of cognitive state detection.

A significant contribution made here is the quantification of behavior from a continuous motor task, in a manner that aligns with the rich psychological literature of flow states (Fig 3A). To that end, we did not detect significant differences between averaged Flow State Survey responses between the task-workload conditions, however, we demonstrate at least modest correlations between individual question ratings, and our novel metric of performance, that suggest this is a quantitative behavioral read out, capable of acting as a proxy for how subjects will rate their flow experience in post-hoc surveys without the need to administer questionnaires and disrupt behavior (Fig 1G versus Fig 3D). Importantly, for any putative deployable product detecting high-performance states, this work has identified what physiological signals are crucial, and in what relative ranking, allowing vendors to align sensor development and inclusion in their platforms accordingly.

## Materials and Methods

### Participants

All protocols and procedures were approved prior to the beginning of study recruitment by the Air Force Research Laboratory Institutional Review Board (AFRL IRB FWR20210142H, spanning August 16^th^ 2021 to August 15^th^ 2026). Study participants (n=19, four female, 29.9 ± 4.8 years) were recruited, verbally briefed on the study protocol and allowed to review an informed consent document before being asked for verbal consent to participate (according to the ethics approved protocol), which was witnessed by laboratory staff whom then noted consent on the copy of the informed consent document provided to the subject and formally enrolled them in the study. Participants were compensated for their time. No minors were included in the current study.

### Data Collection

Upon arrival in the laboratory on the first day participants affirmed their desire to participate in the experiment. They then completed a battery of surveys to capture handedness, demographics, and experience and regularity of video gameplay. A customized version of the shortened Flow State Survey and a Time Dilation survey was administered after the first easy, medium, and hard block trials of each experimental day.

After survey completion, participants are introduced to the game, objectives and controller layout. Emphasis was placed on achieving the highest score by shooting asteroids, with the least amount of ship deaths. Each experimental day, participants first play the baseline calibrating version (∼10 minutes) of the game in the absence of the oddball task. Their performance in this block is used to set their relative asteroid difficulty levels for the rest of the daily session (0.25 x baseline, 1.25 x baseline, and 2 x baseline for E, M and H, respectively). Participants then play through the trial blocks in a pseudo-randomized order (three difficulties x three repeats for nine blocks per day).

Data were collected during gameplay using a BioSemi ActiveTwo electroencephalography system (Cortech Solutions Inc., USA) with 64 channels sampled at 2048 Hz. Peripherals were used to capture participant respiration with a soft belt around the chest, heart rate via clavicle electrodes, ocular artifacts, and galvanic skin conductance on the base of the non-dominant palm of the hand. In several participants galvanic skin response was not recorded due to technical reasons or participant comfort while holding the controller and so it was not analyzed. Participants also wore a commercially available Tobii Pro Glasses 3 (Tobii, Sweden), to capture pupillary activity during the experiment. Game related events and tone presentations were collected by a StimTracker 1G (Cedrus Corporation, USA).

The Air Force Space Shooter task is a custom program, where-in the goal is to both navigate the ship to avoid asteroid collisions and fire the weapon to blow up asteroids for points. Participants used a Logitech G-UF13A controller to play the game and were given a new pair of disposable in-ear headphones each visit.

### Oddball Task

Oddball stimuli were presented randomly but spaced at least 10 and no more than 20 seconds apart. The high pitch pure tone (800 Hz) and the low pitch tone (400 Hz) were presented at a ratio of 1:2. Participants were asked to respond to the tone with their right index finger, by pressing either the upper (high tone) or lower (low tone) bumper buttons on the back of the controller. The in-game sounds were not muted, meaning participants had to hear and distinguish pitch through on-going background noise.

### Data Analysis

All data were analyzed using Python and Python packages. For EEG data the MNE Python package was used (Gramfort et al., 2013). These analyses were run on Lenovo Thinkpad p50 laptops or standard Dell laboratory desktops with Windows operating systems. EEG data were referenced to a common spatial reference, independent component analysis and ocular artifact detection were applied. EEG signals were down-sampled to 256 Hz, and epochs were created from the ten seconds of data prior to and two seconds of data following each pseudo randomly timed tone event. Bad channels throughout a recording were interpolated, if possible, from surrounding electrode sites, then epochs were screened for noise and rejected above 60 µV peak to peak amplitude. The majority of features were extracted from the ten seconds prior to tone onset, but some specifically capturing pupil or EEG peaks after the event were included meaning for most features the identity of the oddball tone was irrelevant. For EEG data and ocular responses these peak and trough detections were set to 0-1s after the oddball tone onset. The final dataset excluded five participant recording days (two days caused by failure in the eye tracking calibration on P1_Day3 and P16_Day1 and three by EEG signal quality P6 day 2, P6 day 3 and P17 day 3). After epoch rejection, total events were 5908 from the original 7560 epochs, from nineteen participants. All EEG power calculations are relative to the other power bands present in the trial (%). Power band bins were 1-4 Hz, 4-8 Hz, 8-12 Hz, 12-25 Hz, 25-45 Hz for delta, theta, alpha, beta, and gamma respectively.

Ocular data were collected on the Tobii Glasses 3 (Tobii, Sweden) system and preprocessed using custom software in SciPy python packages (Virtanen et al., 2020). The Tobii Glasses 3 is a head mounted device capturing pupil diameter and gaze at a 100 Hz sampling rate. The raw pupil diameter data was preprocessed using a multi-stage pipeline to ensure signal quality and reliability for analysis. For each recording, the left and right pupil diameter values were extracted and used to compute the average pupil diameter. The proportion of missing data for the average pupil diameter was quantified and assessed for data completeness. To identify physiologically plausible fluctuations, sample to sample velocity changes in pupil diameter were calculated and any changes exceeding the threshold were marked as invalid and excluded. Additionally, low-value artifacts, typically residual blink samples, were removed using a rolling sample mean deviation threshold. Missing values were interpolated using a linear method applied in both forward and backward directions to preserve temporal continuity. The clean signals were then smoothed using a second-order low pass Butterworth filter with a cutoff frequency of 10 Hz and a sampling rate set at the devices’ native 100 Hz to reduce frequency noise. The processed data was temporally aligned to extract EEG epochs from ten seconds prior to and two seconds following the tone event to extract various features (e.g. pupil diameter std, evoked peak, saccades and sample entropy). All features were calculated in Python packages except saccades which were detected using Tobii’s I-VT (Identification by velocity threshold) algorithm implemented in their proprietary software (Tobii, Sweden).

The ECG and respiration data were recorded simultaneously with EEG using the same BioSemi ActiveTwo system. The ECG data were extracted in MNE and processed in the Heartpy python package (Van Gent et al., 2019). The respiration signal was processed in custom software using band-pass filters to remove baseline drift and high-frequency noise. Peaks and troughs were detected to calculate respiratory cycles. All figures report average and sem unless otherwise noted. ANOVAs were run with the Python statsmodels package using the OLS and Anova_lm methods. Post-hoc tests are pairwise tukey HSDs, except for Fig 2 B, where standard two tailed ttests were used to focus on the within condition comparison, as we were not comparing life lengths between class labels and conditions.

## Acknowledgements

The views expressed are those of the authors and do not reflect the official guidance or position of the United States Government, the Department of Defense, the United States Air Force or the United States Space Force. Appearance of, or reference to, any commercial products or services does not constitute DoD endorsement of those products or services. Imagery in this document are property of the U.S. Air Force. We would like to thank Justin Estepp and William Aue for the support and suggestions during the analyses of these data.

